# Endothelial TMEM16F lipid scramblase regulates angiogenesis

**DOI:** 10.1101/2023.08.17.553724

**Authors:** Ke Zoe Shan, Trieu Le, Pengfei Liang, Ping Dong, Huanghe Yang

## Abstract

Dynamic loss of lipid asymmetry through the activation of TMEM16 Ca^2+^-activated lipid scramblases (CaPLSases) has been increasingly recognized as an essential membrane event in a wide range of physiological and pathological processes, including blood coagulation, microparticle release, bone development, pain sensation, cell-cell fusion, and viral infection. Despite the recent implications of TMEM16F CaPLSase in vascular development and endothelial cell-mediated coagulation, its signaling role in endothelial biology remains to be established. Here, we show that endothelial TMEM16F regulates *in vitro* and *in vivo* angiogenesis through intracellular signaling. Developmental retinal angiogenesis is significantly impaired in TMEM16F deficient mice, as evidenced by fewer vascular loops and larger loop areas. Consistent with our *in vivo* observation, TMEM16F siRNA knockdown in human umbilical vein endothelial cells compromises angiogenesis *in vitro*. We further discovered that TMEM16F knockdown enhances VE-cadherin phosphorylation and reduces its expression. Moreover, TMEM16F knockdown also promotes Src kinase phosphorylation at tyrosine 416, which may be responsible for downregulating VE-cadherin expression. Our study thus uncovers a new biological function of TMEM16F in angiogenesis and provides a potential mechanism for how the CaPLSase regulates angiogenesis through intracellular signaling.

## Introduction

Angiogenesis, the formation of new blood vessels, is indispensable for development, growth, and wound healing (Carmeliet and Jain, 2011; Chung, Lee and Ferrara, 2010). Aberrant angiogenesis, on the other hand, fuels the pathogenesis of cancer, inflammatory diseases, hypertension, and ocular disorders (Carmeliet and Jain, 2011; Chung, Lee and Ferrara, 2010). A major angiogenesis regulator in endothelial cells (ECs) is VEGF signaling, which regulates EC proliferation, migration, adhesion, and survival through phospholipase Cγ, nuclear factor of activated T cells (NFAT), and Src tyrosine kinase pathways (Simons, Gordon and Claesson-Welsh, 2016). While numerous anti-angiogenic agents, such as VEGF-targeted drugs, have been developed to mitigate pathological angiogenesis-associated diseases, these agents have limited efficacy and show transient therapeutic effects (Eelen et al., 2020; Jaszai and Schmidt, 2019). Thus, it is urgent to identify new molecular players in angiogenic pathways to treat cancer, ocular, and other angiogenesis-associated diseases that have caused global health burdens.

Recent studies show that the TMEM16F deficiency impairs blood vessel development in the murine placentas (Zhang et al., 2020a) or the zebrafish embryos (Delcourt et al., 2015). However, it is unclear whether the observed defects are directly derived from TMEM16F deficiency in ECs and how TMEM16F regulates vascular development. TMEM16F is a CaPLSase that rapidly collapses membrane phospholipid asymmetry by bi-directionally and non-selectively transporting all major phospholipids down their concentration gradients, leading to surface exposure of phosphatidylserine (PS) (Bevers and Williamson, 2016; Kunzelmann et al., 2014; Suzuki et al., 2010). Despite of the growing list of its functions in blood coagulation (Bevers et al., 1992; Fujii et al., 2015; Schmaier et al., 2023; Yang et al., 2012; Yu et al., 2021), cell fusion (Braga et al., 2021; Zhang et al., 2020a), plasma membrane repair (Wu et al., 2020), bone development(Ehlen et al., 2013; Ousingsawat et al., 2015), viral infection (Zaitseva et al., 2017), and neuron maintenance (Cui et al., 2023; Soulard et al., 2020; Zhang et al., 2020b), the role of TMEM16F in endothelial biology just started emerging (Schmaier et al., 2023; Yu et al., 2021). Here we report that TMEM16F CaPLSase is functionally expressed in ECs and plays an important role in regulating neonatal retinal angiogenesis *in vivo* and human umbilical vein endothelial cell (HUVEC) angiogenesis *in vitro*. TMEM16F deficiency in HUVECs enhances Src tyrosine kinase activity, which leads to phosphorylation and downregulation of VE-cadherin and subsequent defects in angiogenesis *in vitro*. Our findings thus establish a physiological function of TMEM16F in angiogenesis and reveal the Src signaling pathway as a downstream target of TMEM16F.

## Results and Discussion

### TMEM16F is a major CaPLSase in endothelial cell lines

In order to understand TMEM16F’s physiological function in ECs, it is critical to quantitatively measure its contribution to Ca^2+^-activated lipid scrambling. To achieve this, we first used siRNAs to silence *TMEM16F* in two widely used primary human EC models, HUVECs and human aortic endothelial cells (HAECs). Our western blot shows that the siRNA knockdown nearly completely eliminates TMEM16F protein expression in both EC cell lines (Figs. 1A-B and S1A-B). Next, we employed the fluorescence imaging-based assay (Le et al., 2019; Le, Le and Yang, 2019) to quantify CaPLSase activity in the control siRNA and TMEM16F siRNA transfected endothelial cell lines (Fig. 1C). In the control siRNA-transfected HUVECs and HAECs, Ca^2+^ ionophore ionomycin triggers robust PS exposure on EC surfaces, which is reported by the time-dependent accumulation of fluorescently conjugated Annexin V (AnV), a PS-binding protein (Figs.1 D-F and S1C-E). In contrast, PS exposure kinetics and maximum exposure levels were dramatically diminished by TMEM16F knockdown (Figs.1D-F and S1C-E). To further confirm our findings using western blot and the lipid scrambling assay, we conducted a whole-cell patch clamp to measure TMEM16F-mediated current in HUVECs, utilizing its moonlighting function as a Ca^2+^-activated non-selective ion channel (Le et al., 2019; Liang and Yang, 2021; Yang et al., 2012; Yu et al., 2015). We found that the typical Ca^2+^- and voltage-activated, outward rectifying TMEM16F current (Yang et al., 2012) was largely diminished from the TMEM16F siRNA transfected HUVECs (Fig. 1G-H). Taken together, our quantitative analysis using three different approaches establishes that TMEM16F is functionally expressed in ECs and is a major EC CaPLSase.

**Figure 1.**
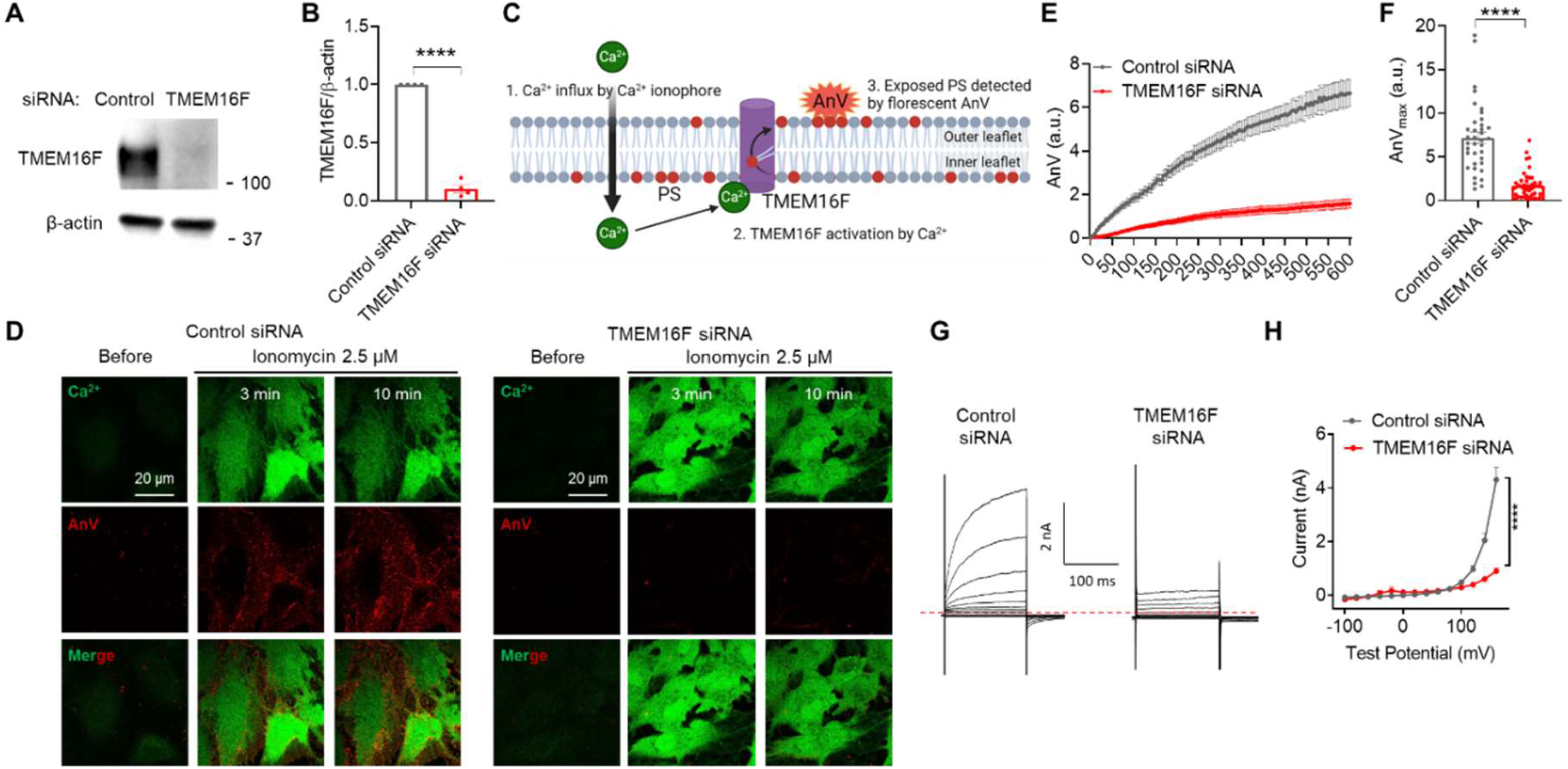
TMEM16F is functionally expressed in endothelial cells. (A) Representative western blots of TMEM16F in HUVECs with and without siRNA knockdown. (B) Densitometry quantifications of TMEM16F and loading control β-actin (n=4). (C) Schematic representation of fluorescence-based scrambling assay. PS: phosphatidylserine, AnV: Annexin V. (D) Representative images of Ca^2+^ and AnV in control (left) and TMEM16F knockdown (right) HUVECs stimulated with 2.5 μM ionomycin. (E, F) Quantifications of the time course or the maximum fluorescence intensity of AnV at 10 minutes post ionomycin stimulation for control siRNA (n=37) and TMEM16F (n=46). Each dot represents AnV signals from one cell. Data are presented as mean±S.E.M. ****P<0.0001, two-tailed t-test. (G) Representative currents recorded in control siRNA- and TMEM16F siRNA-treated HUVECs. The currents were elicited by a voltage step protocol from -100 mV to +160 mV with a 20 mV increment. Holding potential was set at -60 mV. (H) Current-voltage (*I-V*) relationship of currents recorded in (G). Data are presented as mean±S.E.M. Currents at +160 mV were compared with two-tailed t-test ****P<0.0001 (n=7 for each group).

### TMEM16F deficiency leads to defective retinal angiogenesis in mice

The angiogenic development of the mouse retina has been extensively used to evaluate angiogenesis defects *in vivo (Stahl et al*., *2010)*. To evaluate TMEM16F’s angiogenic role *in vivo*, we quantified the vascular network in the retinas from postnatal day 5 (P5) wildtype (WT) and TMEM16F knockout (KO) mice in C57Bl/6J background. Our IB4 staining of the whole mount retinas showed that the vasculature coverage of TMEM16F KO mice is significantly reduced compared to that of their WT littermates (Fig. 2A-B top panels and 2C). Our unbiased analysis of retinal vasculature complexity (Fig. 2A-B middle and bottom panels, see Methods) further demonstrates that the average number of vascular loops per field of view is markedly lower (Fig. 2D) and the average hole areas of the vascular loop are significantly higher (Fig. 2E) in TMEM16F KO retinas. Our analysis of the retinal developmental angiogenesis of TMEM16F KO mice confirms that TMEM16F plays an important role in angiogenesis *in vivo*.

**Figure 2.**
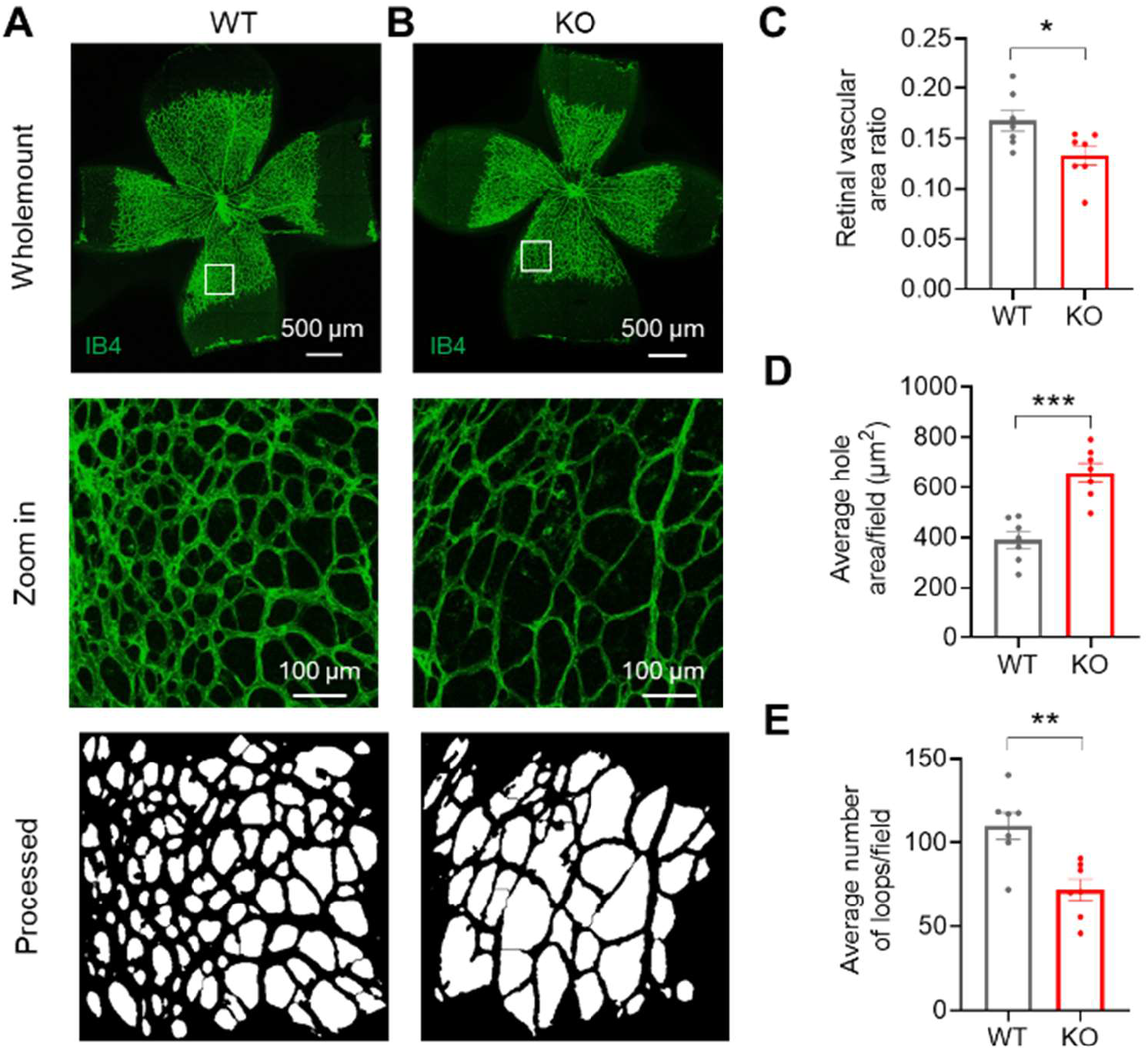
TMEM16F deficiency suppresses angiogenesis *in vivo*. (A-B) IB4 (green) staining of retinal whole mount from P5 WT (A) and TMEM16F KO (B) littermate. White boxes indicate enlarged area shown in the middle panels. The bottom panels illustrate processed enlarged area used for the calculations of number of loops and hole areas. White areas are defined as holes and vasculature (black lines) encircling the holes as loops. (C-E) Quantification of retinal vasculature coverage (C), average hole areas (D), and number of loops (E) calculated from four random fields of each retina. Data are presented as mean±S.E.M. Each dot represents data from one animal (n=7 pairs of WT and TMEM16F KO littermates). *P<0.05, **P<0.01, ***P<0.001, two-tailed t-test.

### TMEM16F deficiency in ECs compromises angiogenesis *in vitro*

To assess the endothelial-specific role of TMEM16F in angiogenesis, we examined the impact of TMEM16F siRNA on HUVEC tube formation *in vitro*. Our time-lapse imaging reveals that the HUVECs transfected with control siRNA successfully form a robust, three-dimensional network of tube-like structures on the Matrigel matrix within four hours of seeding (Fig. 3A and Video S1). Following this initial formation, the network undergoes gradual remodeling. This process entails the subtle merging of proximate tubes, leading to the creation of larger loop-like structures. 24 hours post seeding, there are 4.56 ± 0.44 loops/mm^2^ remained in control siRNA transfected HUVECs (Fig. 3B). In stark contrast, the tube network formed by the TMEM16F siRNA transfected HUVECs is dramatically unstable as evidenced by the large loops and significantly decreased loop density (1.74 ± 0.13 loops/mm^2^) 24 hours post seeding (Fig. 3 and Video S1). Close examination of the tube network at different time points reveals that the tubes formed by the TMEM16F deficient cells start to break apart about 8 hours post seeding, and the separated cells tend to aggregate into large cell clusters at tube junctions, which results in markedly reduced loop density. Consistent with the *in vivo* angiogenesis defects in TMEM16F KO mice (Fig. 2), our *in vitro* angiogenesis experiments further support that endothelial TMEM16F plays an important role in the development of blood vessels.

**Figure 3.**
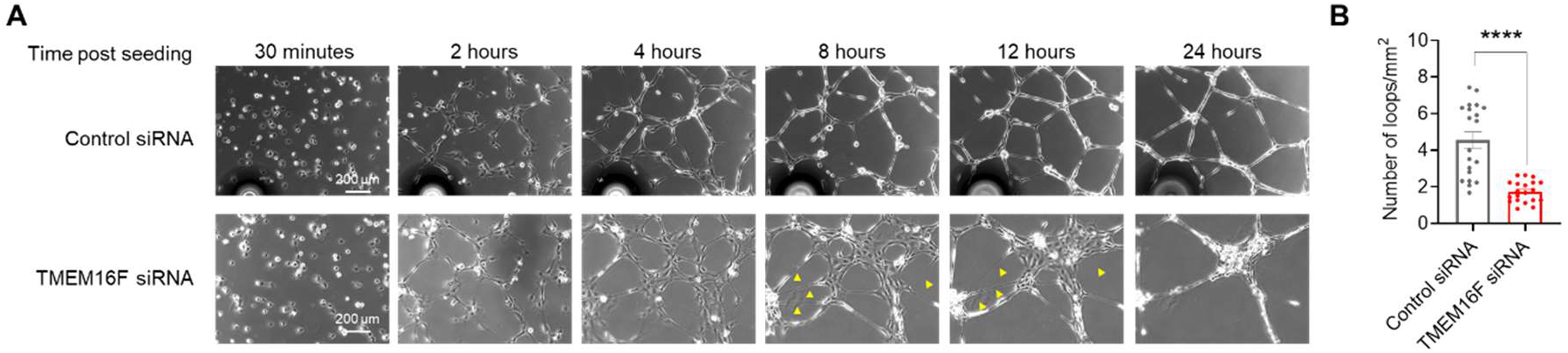
TMEM16F deficiency in HUVEC impairs *in vitro* angiogenesis. (A) Representative images of tube-forming control and TMEM16F knockdown HUVECs after seeded on Matrigel. Yellow arrowheads indicate the thin and stretched tubes. (B) Quantification of loop numbers of control (n=20) and TMEM16F (n=20) knockdown HUVECs at 24 hours after seeding. Each dot represents data from one well of tube formation μ-Slide. Data are presented as mean±S.E.M. ****P<0.0001, two-tailed t-test.

### TMEM16F deficiency downregulates VE-cadherin by promoting Src and VE-cadherin phosphorylation

The instability of the tubes formed by the TMEM16F knockdown HUVECs (Fig. 3) suggests that the EC cell-cell junctions may be impaired by TMEM16F deficiency. The weakened junctions may fail to hold the tubes together (Fig. 3, 8-24 hours), leading to unstable tube networks. To assess whether TMEM16F regulates the adherens junctions of ECs, we measured the expression of VE-cadherin, an endothelial adhesion protein that is required at the cell-cell junctions to stabilized and maintain the newly-formed vessels(Bentley et al., 2014). Our western blot results reveal that TMEM16F siRNA knockdown significantly downregulates VE-cadherin expression (Fig. 4A-B). A prominent pathway destabilizing VE-cadherin junctions is Src kinase phosphorylation at tyrosine (Tyr) 685 of VE-cadherin, causing VE-cadherin endocytosis (Vestweber, 2008; Wallez et al., 2007), which could lead to degradation(Vincent et al., 2004). To evaluate whether TMEM16F deficiency in HUVECs increases VE-cadherin phosphorylation by Src kinase, we examined VE-cadherin phosphorylation levels. Our western blot results indicate that TMEM16F knockdown in HUVECs robustly increased VE-cadherin phosphorylation at Tyr 685 (Fig. 4C-D), suggesting that Src activity is upregulated. Indeed, the active form of Src kinase, pY416-Src, is significantly enhanced in TMEM16F knockdown HUVECs (Fig. 4E-F). Taken together, our biochemical evidence indicates that TMEM16F deficiency in ECs promotes Src kinase phosphorylation and activation, which phosphorylates VE-cadherin, and could subsequently lead to VE-cadherin endocytosis and degradation (Fig. 4G). Downregulation of VE-cadherin likely destabilizes the EC junctions, resulting in impaired stability of newly formed blood vessels and the observed angiogenesis defects.

**Figure 4.**
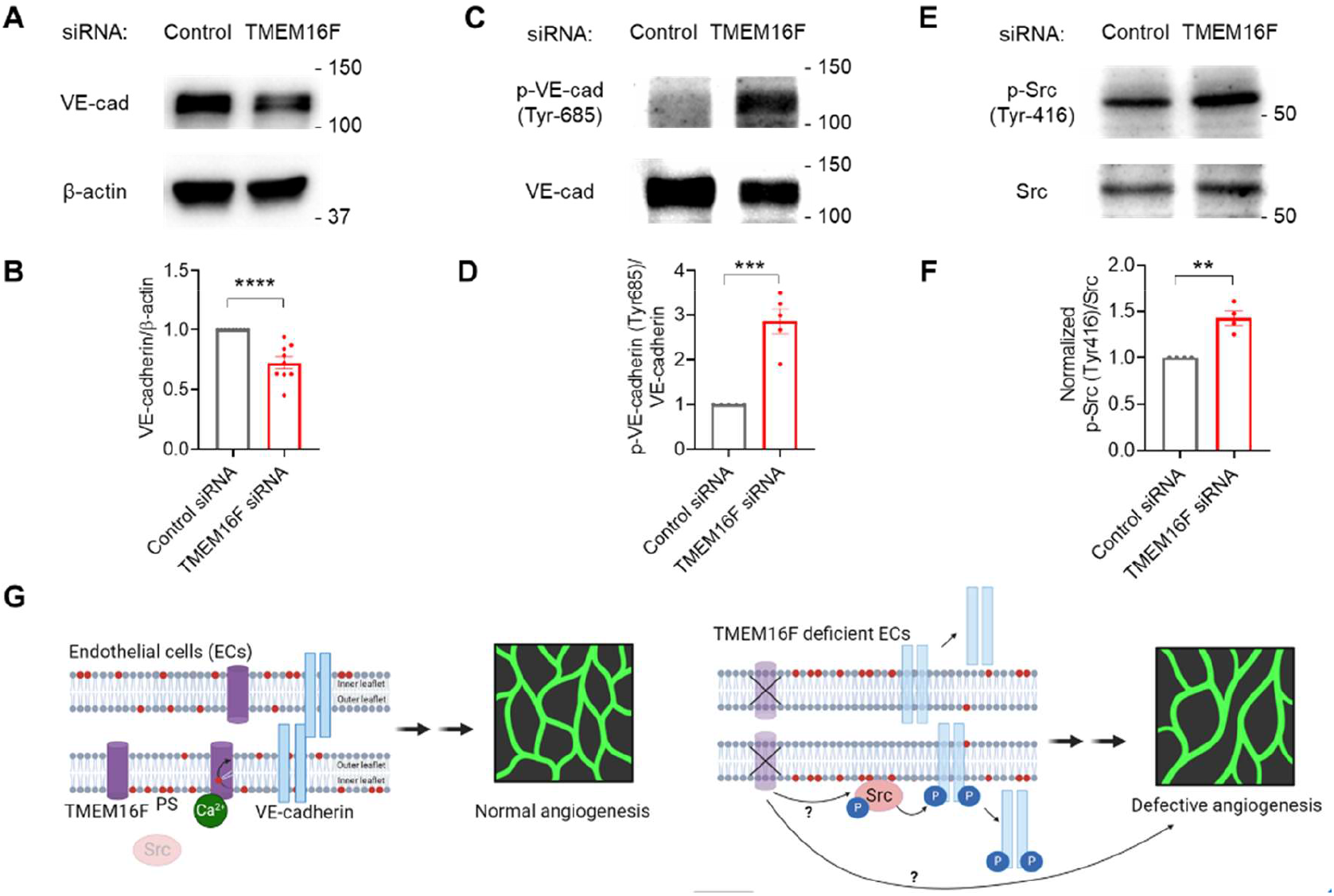
TMEM16F knockdown reduces VE-cadherin expression and promotes the phosphorylation of VE-cadherin and Src. (A,B) Representative western blot of anti-VE-cadherin (A) and densitometry quantifications (n=9) (B). (C,D) Representative western blot of anti-Tyr685 phosphorylated and anti-total VE-cadherin (C) and densitometry quantifications (n=5) (D). (E,F) Representative western blot anti-Tyr416 phosphorylated and anti-total Src (E) and densitometry quantifications (n=4) (F). (G) Proposed role of TMEM16F scramblase in angiogenesis. TMEM16F deficiency increases Src phosphorylation and promotes its activation, which downregulates VE-cadherin and leads to defective angiogenesis. Data are presented as mean±S.E.M. **P<0.01, ***P<0.001, ****P<0.0001, two-tailed t-test.

In summary, utilizing multidisciplinary approaches including *in vivo* mouse retina model, *in vitro* angiogenesis model, western blot, fluorescence imaging, and patch-clamp recording, we show that TMEM16F deficiency impairs *in vivo* mouse retina angiogenesis and *in vitro* tube formation, and alters Src-VE-cadherin signaling in ECs (Fig. 4G). Our findings offer insights that could pave the way for a deeper understanding of lipid scramblase-driven cell signaling. This knowledge may aid in developing precision therapeutics targeting CaPLSase in ECs to treat diseases dependent on angiogenesis.

Current research on lipid scramblases has been primarily focused on their activity to promote PS exposure and subsequent extracellular signaling that requires surface-exposed PS, such as the recruitment of PS-binding clotting factors for blood coagulation (Schmaier et al., 2023; Yu et al., 2021), PS receptors for phagocytosis(Zhang et al., 2020b), the ADAM proteases for proteolytic cleavage (Bleibaum et al., 2019; Sommer et al., 2016), and viral envelop proteins for infection (Zaitseva et al., 2017). In this study, however, we uncover that Src kinase and VE-cadherin are novel intracellular signaling targets of TMEM16F in ECs (Fig. 4G). Although the exact underlying mechanism warrants future investigation, TMEM16F deficiency-induced VE-cadherin downregulation and the upregulation of phosphorylated Src kinase and VE-cadherin set a clear example that TMEM16F CaPLSase-mediated loss of phospholipid asymmetry on the plasma membrane can alter intracellular signaling.

Interestingly, the loss of TMEM30A, a critical regulatory subunit of phospholipid flippases that help maintain membrane lipid asymmetry (Bryde et al., 2010; Nagata, Sakuragi and Segawa, 2020), inhibits Src kinase phosphorylation in ECs (Zhang et al., 2019). Different from scramblase deficiency that makes it difficult to lose lipid asymmetry, the deficiency of the ATP-driven flippases leads to loss of lipid asymmetry and surface exposure of PS and phosphatidylethanolamine (Bevers and Williamson, 2016; Nagata, Sakuragi and Segawa, 2020). The opposite effects of flippase deficiency and TMEM16F scramblase deficiency on Src kinase phosphorylation in ECs suggest that the dynamic transbilayer redistribution of phospholipids mediated by these lipid transporters can indeed trigger intracellular changes. One such possible regulatory mechanism could be the membrane recruitment of Src kinases, a prerequisite for their functions (Shvartsman et al., 2007). To be tightly associated with the plasma membrane, Src kinase preferentially bind to negatively-charged phospholipids such as PS through a stretch of positively-charged residues at the N-terminus (Murray et al., 1998; Sigal et al., 1994) (Fig. 4G). Indeed, recent studies show that scramblase-mediated membrane lipid redistribution disrupts the membrane association of Src kinases, resulting in their translocation from the plasma membrane to the cytosol (Wu, Song and Veillette, 2021; Yeung et al., 2006), albeit the molecular identities of the lipid scramblases remain elusive.

A recent report suggests that in addition to TMEM16F, another TMEM16 CaPLSase (Whitlock et al., 2018), TMEM16E also expresses in ECs (Schmaier et al., 2023). Ca^2+^ influx triggers PS exposure through TMEM16F and TMEM16E, which promotes procoagulant activity on the EC surface. Intriguingly, the double knockdown of TMEM16F and TMEM16E does not lead to an additive reduction of procoagulant activity, and the silencing of either TMEM16F or TMEM16E eliminates most of the Ca^2+^-induced PS exposure, suggesting that the two scramblases could be epistatic to one another (Schmaier et al., 2023). TMEM16E likely compensates for and counterbalance the loss of TMEM16F *in vivo*. This may explain why the retinal angiogenesis is not completely inhibited in TMEM16F knockout mice. Future *in vitro* and *in vivo* studies using the double knockout of endothelial TMEM16F and TMEM16E are required to further dissect the roles of the CaPLSases in vessel development and other endothelial functions.

Dysregulation of angiogenesis is a hallmark of highly prevalent diseases such as cancer, cardiovascular and inflammatory diseases, and ocular disorders (Carmeliet and Jain, 2000). However, the success of anti-angiogenesis therapies, which predominantly target pro-angiogenic factors, is curtailed by the activation of alternative angiogenic pathways and resistance mechanisms (Eelen et al., 2020). To overcome these limitations, it is critical to further understand angiogenesis mechanisms and discover new pro-angiogenic pathways. Interestingly, PS exposure on the endothelium is frequently observed in conditions like cancer (Ran, Downes and Thorpe, 2002; Zhang et al., 2017), inflammatory bowel disease (Zhang et al., 2021), and choroidal neovascularization (Li et al., 2015). Studies have shown that anti-PS antibodies or PS-binding proteins can inhibit vessel formation (Li et al., 2015; Ran, Downes and Thorpe, 2002; Zhang et al., 2017; Zhang et al., 2021). Therefore, targeting PS exposure in ECs presents a new strategy for designing more effective anti-angiogenesis therapies. Identifying TMEM16F as a primary EC CaPLSase that mediates PS exposure and understanding its role in regulating Src-VE cadherin signaling, open a new avenue to specifically target lipid scramblases for treating angiogenesis-dependent diseases.

## Methods Mice

The TMEM16F knockout (KO) mouse line was a gift from Dr. Lily Jan (UCSF) and has been reported previously (Yang et al., 2012; Zhang et al., 2020a). The heterozygous mice have been backcrossed ten generations into a C57BL6/J strain and were used to generate TMEM16F^−/−^ KO mice. Mouse handling and usage were carried out in strict compliance with protocols approved by the Institutional Animal Care and Use Committee at Duke University in accordance with National Institutes of Health guidelines.

### Retinal flatmount and staining

Enucleated eyes from wildtype or TMEM16F KO littermate pairs at postnatal day 5 were fixed in 4% paraformaldehyde in phosphate-buffered saline (PBS) overnight at 4°C with gentle shaking. Retinas were isolated from enucleated eyes and were blocked and permeabilized with 1% BSA and 0.5 % Triton X-100 in PBS overnight at 4°C. The retinal vasculatures were then stained with Fluorescein-conjugated isolectin B4 (IB4, 1:100 dilution, Vector laboratories, #FL-1201) overnight at 4°C with gentle shaking. After PBS washing, the retinas were flat-mounted and imaged with Zeiss 780 inverted confocal microscope. The area of retinal vasculature holes and the number of loops per field of view were evaluated. For each retina, four square subregions were randomly chosen by an investigator who was unaware of the retina’s genotype. We utilized a custom MATLAB script (available at https://github.com/superdongping/Retina_blood_vessel) to measure the vascular area, hole area, and loop count in these subregions, leveraging the built-in function ‘imbinarize’. Processed samples of the vascular area can be viewed in Fig. 2.

### Cell culture and siRNA transfection

HUVECs and HAECs (obtained from Lonza and authenticated by the Duke Cell Culture Facility) were cultured in EGM-2 BulletKit (Lonza, #CC-3162) in a humidified incubator at 37°C and 5% CO_2_-95% air. For imaging assays, HUVECs and HAECs cells were seeded on coverslips coated with poly-L-lysine (Sigma-Aldrich, #P2636) in 24-well plates. For western blot, HUVECs and HAECs cells were seeded in 10-cm cell culture dishes. For siRNA transfection, TMEM16F was knocked down by transfecting with non-targeting Silencer Negative Control No. 2 siRNA (Invitrogen, #AM4637) or TMEM16F SMARTpool siRNA (a pre-mixed pool of four siRNAs from Dharmacon Research, #M-003867-01-0020) using Lipofectamine RNAiMAX Transfection Reagent (Invitrogen, #13778075) following the manufacturer’s instructions (volume ratio-RNAiMAX: 10 μM siRNA= 2: 1). Fresh medium was changed the next day, and cells were cultured for another 24 hours before conducting other assays.

### Fluorescence imaging of Ca^2+^ and PS exposure

This assay was done as reported previously (Fig. 1C) (Le et al., 2019; Le, Le and Yang, 2019). Briefly, Calbryte 520 AM (AAT Bioquest, #20701) was used to monitor Ca^2+^ dynamics, and CF 594-tagged AnV (Biotium, #29011) was used to detect exposed PS. ECs were stained with 1 μM Calbryte 520 AM for 10 minutes before transferring to AnV solution (1:150 in Hank’s balanced salt solution, Gibco, #14025-092). Time-lapse imaging was taken before and after 2.5 μM ionomycin stimulation with Zeiss 780 inverted confocal microscope to monitor Ca^2+^ dynamics and AnV binding. A custom MATLAB code (available at https://github.com/YZ299/matlabcode/blob/main/matlabcode.m) used to quantify the PS signal intensities was reported previously (Le et al., 2019; Le, Le and Yang, 2019).

### *In vitro* tube formation assays

Prior to cell seeding, 10 μl of Matrigel (Corning, # 356231, lot #2213003) was added to each well of the µ-Slide 15 Well 3D (ibidi, #81506) without touching the wall of the wells to avoid unlevel matrix formation. Matrigel was then allowed to polymerize by incubating the µ-Slide at 37°C for 45 minutes to 1 hour. After Matrigel solidified, 50 μl of 5.5×10^5^ cells/ml HUVECs in EGM2 were added to each well. For time-lapse imaging, images were captured 30 minutes after seeding at a 10-minute interval using the Zeiss Axio Observer Z1 microscope. For loop numbers quantification, µ-Slides were imaged 24 hours after seeding using an Olympus IX71 inverted epi-fluorescent microscope (Olympus IX73) with a 4x objective. Loop numbers were quantified from the whole well with ImageJ and calculated as loop numbers/mm^2^ by dividing the µ-Slide inner well growth area.

### Immunoblotting

After trypsinization and PBS washing, HUVECs with control or TMEM16F siRNA knockdown were harvested in lysis radioimmunoprecipitation assay (RIPA) lysate buffer (Thermo Fisher Scientific, #89900) supplemented with 1X Halt™ Protease and Phosphatase Inhibitor Cocktail (Thermo Fisher Scientific, #78440). 20-30 μg of proteins were mixed with 1× Laemmli (Bio-Rad, #161-0747) and 2-mercaptoethanol and loaded onto SDS– polyacrylamide gel electrophoresis (PAGE) gels. After separation, the Trans-Blot Turbo Transfer system (Bio-Rad) was used to transfer the proteins from the SDS-PAGE gels to polyvinylidene difluoride membranes. The membranes were blocked with 5% bovine serum albumin (BSA, Tocris, #5217) in TBST (TBS supplemented with 0.1% Tween 20) at room temperature for at least an hour. Following blocking, membranes were incubated in anti-VE-cadherin (1:1000 dilution, Santa Cruz, #sc-9989), anti-phospho-VE-cadherin (Tyr685) (1:1000 dilution, ECM bioscience, #CP1981), anti-Src (1:1000 dilution, ABclonal, #A11707), anti-phospho-Src (Tyr-416) (1:1000 dilution, ABclonal, #AP0452), or β-actin (1:10000 dilution, Sigma, #A1978) overnight at 4°C. Membranes were washed with TBST the next day and incubated with HRP-conjugated goat anti-rabbit-IgG secondary antibody (Sigma, #A0545) or anti-mouse-IgG secondary antibody (Sigma, #A1917) for 1 hour at room temperature before detecting with the Clarity Western ECL Substrate Kit (Bio-Rad, #170-5060).

### Electrophysiology

TMEM16F currents were recorded in whole-cell configuration using an Axopatch 200B amplifier (Molecular Devices) and the pClamp software package (Molecular Devices). Glass pipettes were pulled from borosilicate capillaries (Sutter Instruments) and fire-polished using a microforge (Narishge) to reach a resistance of 2–3 MΩ. The pipette solution (internal) contained (in mM) 140 CsCl, 1 MgCl_2_, and 10 HEPES, plus 1 CaCl_2_ to avoid the long-delay activation of TMEM16F(Liang and Yang, 2021; Zhang et al., 2022). The bath solution contained 140 CsCl, 10 HEPES, and 5 EGTA. Adjusted to pH 7.3 for both sides with CsOH. Currents were recorded about 2 minutes after whole-cell formation to ensure the activation of TMEM16F(Liang and Yang, 2021; Zhang et al., 2022).

Currents were normalized to cell capacitance as current density. *I-V* curves were constructed from steady-state currents after depolarization. Individual *I-V* curves were fitted with a Boltzmann function,

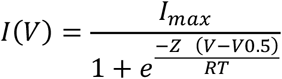

where *I*_*max*_ denotes the maximal current, *V*_*0*.*5*_ denotes the voltage of half-maximal activation of conductance, *z* denotes the net charge moved across the membrane during the transition from the closed to the open state, and *F* denotes the Faraday constant.

### Statistical analysis

All statistical analyses were performed in Prism software (GraphPad). Unpaired two-tailed Student’s t-test was used for single comparisons between two groups, and one-way ANOVA (with Tukey’s multiple comparisons test) was used for multiple comparisons. Sample number (n) values are indicated in the results section and figure legends. All data are presented as the mean ± standard error of the mean (S.E.M.). Symbols *, **, ***, ****, and ns denote statistical significance corresponding to p-value<0.05, <0.01, <0.001, <0.0001, and no significance, respectively.

## Acknowledgments

The authors thank Timothy McCord, Xi Chen, and Christopher Kontos for their help during the course of this work. This work was supported by the NIH-DP2GM126898 grant (awarded to H.Y.) and American Heart Association Predoctoral fellowship (#20PRE35120162, awarded to T.L.).

## Author contributions

Conceptualization: H.Y., T.L., K.Z.S.; Methodology: K.Z.S., T.L., P.L., P.D., H.Y.; Investigation: K.Z.S., T.L., P.L., Formal analysis: K.Z.S., T.L., P.L., P.D.; Validation: K.Z.S.; Visualization: K.Z.S.; Writing - original draft: K.Z.S., H.Y.; Resources: H.Y., Supervision: H.Y.; Funding acquisition: H.Y.; Project administration: H.Y.

## SUPPLEMENTARY FILE

**Figure S1.**
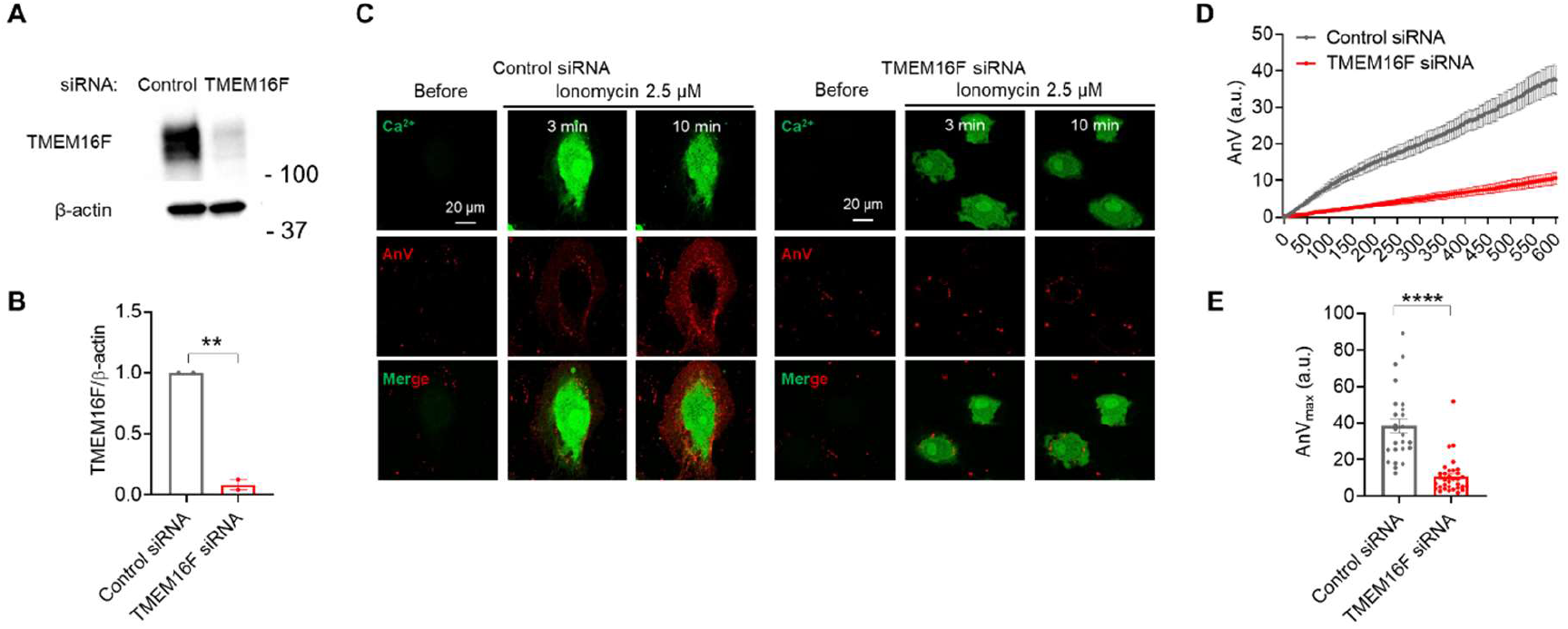
TMEM16F is functionally expressed in human aortic endothelial cells (HAECs). (A) Representative western blot of TMEM16F in HAECs with and without siRNA knockdown. (B) Densitometry quantifications of TMEM16F and loading control β-actin (n=2). (C) Representative images of Ca^2+^ and AnV in control (left) and TMEM16F knockdown (right) HAECs stimulated with 2.5 μM ionomycin. (D,E) Quantifications of the time course (D) or the maximum fluorescence intensity of AnV 10 minutes post ionomycin stimulation (E) for control siRNA (n=25) and TMEM16F (n=33). Each dot represents PS signals from one cell. Data are presented as mean±S.E.M. **P<0.01, ****P<0.0001, two-tailed t-test.

**Movie S1. TMEM16F deficiency reduces tube formation**.

Tube-forming control and TMEM16F knockdown HUVECs after seeded on Matrigel. Time stamps (hour: minute: second) shown are post-seeding time.

